# The Gene Expression Profile of Uropathogenic *Escherichia coli* in Women with Uncomplicated Urinary Tract Infections Is Recapitulated in the Mouse Model

**DOI:** 10.1101/2020.02.18.954842

**Authors:** Arwen E. Frick-Cheng, Anna Sintsova, Sara N. Smith, Michael Krauthammer, Kathryn A. Eaton, Harry L. T. Mobley

## Abstract

Uropathogenic *Escherichia coli* (UPEC) is the primary causative agent of uncomplicated urinary tract infections (UTIs). UPEC fitness and virulence determinants have been evaluated in a variety of laboratory settings that include a well-established mouse model of UTI. However, the extent to which bacterial physiology differs between experimental models and human infections remains largely understudied. To address this important question, we compared the transcriptomes of three different UPEC isolates in human infection and a variety of laboratory conditions including LB culture, filter-sterilized urine culture, and the UTI mouse model. We observed high correlation in gene expression between the mouse model and human infection in all three strains examined (Pearson correlation coefficient of 0.86-0.87). Only 175 of 3,266 (5.4%) genes shared by all three strains had significantly different expression levels, with the majority of them (145 genes) down-regulated in patients. Importantly, gene expression of both canonical virulence factors and metabolic machinery were highly similar between the mouse model and human infection, while the *in vitro* conditions displayed more substantial differences. Interestingly, comparison of gene expression between the mouse model and human infection hint at differences in bladder oxygenation as well as nutrient composition. In summary, our work strongly validates the continued use of this mouse model for the study of the pathogenesis of human UTI.

**Importance:** Different experimental models have been used to study UPEC pathogenesis including *in vitro* cultures in different media, tissue culture, as well as mouse models of infection. The latter is especially important since it allows evaluation of mechanisms of pathogenesis and potential therapeutic strategies against UPEC. Bacterial physiology is greatly shaped by environment and it is therefore critical to understand how closely bacterial physiology in any experimental model relates to human infection. In this study, we found a very strong correlation in bacterial gene expression between the mouse model and human UTI using identical strains, suggesting that the mouse model accurately mimics human infection, definitively supporting its continued use in UTI research.

## Introduction

Urinary tract infections (UTIs) are one of the most common bacterial infections in otherwise healthy individuals. Over 50% of women will experience at least one UTI in their lifetime, and half of these women will experience a recurrent infection within a year (1, 2). These infections affect 150 million people per year and result in annual medical costs of $3.5 billion in the US alone (3). Uropathogenic *Escherichia coli* (UPEC) is responsible for 80% of uncomplicated UTIs (1) and deploy diverse strategies to survive and replicate in the human host. These comprise an array of virulence factors including, but not limited to, iron acquisition systems (siderophores and heme receptors), fimbriae and other adhesins, flagella, and toxins (4-7). The importance of these systems to bacterial fitness has been studied in detail using multiple models including cultures in laboratory media, human urine cultures, tissue culture, and a mouse model first established over 30 years ago (8). However, animal models can fail to recapitulate important aspects of the human response to disease (9). Whether the mouse model accurately reflects the native environment found during human infection has not been adequately addressed. Therefore, it is vitally important to determine if the mouse model of ascending UTI recapitulates human UTI since defining mechanisms of pathogenesis and the development of UTI therapies relies on this assumption (10).

Previous studies compared the mouse model to human UTIs using microarrays to assess differences in bacterial gene expression (11, 12). Initially, urine from mice infected with UPEC type strain CFT073 was collected over a period of ten days, pooled and analyzed using a microarray based on the CFT073 genome (11). In a follow-up study, urine was collected from eight women with complicated UTIs and bacterial gene expression in the human host was analyzed, again using microarrays based on the CFT073 genome (12). Relative expression levels of 46 fitness genes were compared between the mouse model and human UTI. This comparison demonstrated a Pearson’s correlation coefficient of 0.59 and was strongest for iron acquisition systems and weakest for adhesin and motility systems (12). While encouraging, this study did not provide conclusive evidence that the mouse model closely replicated human UTI. A key weakness of the previous comparison was that genetic differences between currently circulating isolates and strain CFT073 used for the mouse infections would obscure strain-specific responses, either due to differences in mouse *versus* human UTI or because the CFT073-specific microarrays would not detect expression of genes that are not encoded by that strain.

We have recently used RNA sequencing (RNA-seq) to quantify the UPEC transcriptome during acute infection in 14 female patients (13). Importantly, RNA-seq is a more comprehensive platform to analyze the transcriptome of clinical UPEC strains since, unlike microarrays, it is not limited by strain-specific probes. In this study, we report the transcriptome during murine UTI for three of the 14 clinical strains using RNA-seq and directly compare the gene expression patterns for these identical strains between human UTI and the mouse model. We observed a high correlation between human infection and mouse infection (Pearson correlation coefficient ranging from 0.86-0.87) with only 175 of 3,266 shared genes being differentially expressed. Gene expression of classical virulence factors as well as metabolic genes in the mouse model closely resembled those observed during human UTI. Our study is the first of its kind to directly compare the bacterial transcriptomes between human and mouse UTI using identical strains. We conclude that the mouse model accurately reflects bacterial gene expression observed during human infection.

## Results

### Study Design

We previously sequenced the transcriptomes of 14 UPEC strains isolated directly from the urine of patients with uncomplicated UTIs (hUTI) and immediately stabilized with RNAprotect (13). Three out of 14 strains (HM43, HM56, and HM86) were chosen to conduct transcriptomic studies in the prevailing mouse model of UTI (mUTI) (8). We selected strains whose hUTI transcriptomes had the highest proportion of bacterial reads to eukaryotic reads (**Supplemental Table 1**) and that possessed a prototypical UPEC virulence factor profile. All three strains belong to the B2 phylogroup (13), where the majority of UPEC strains reside, and encode a range of siderophores, heme receptors, as well as multiple fimbrial types (**Fig. 1**).

**Fig. 1.**
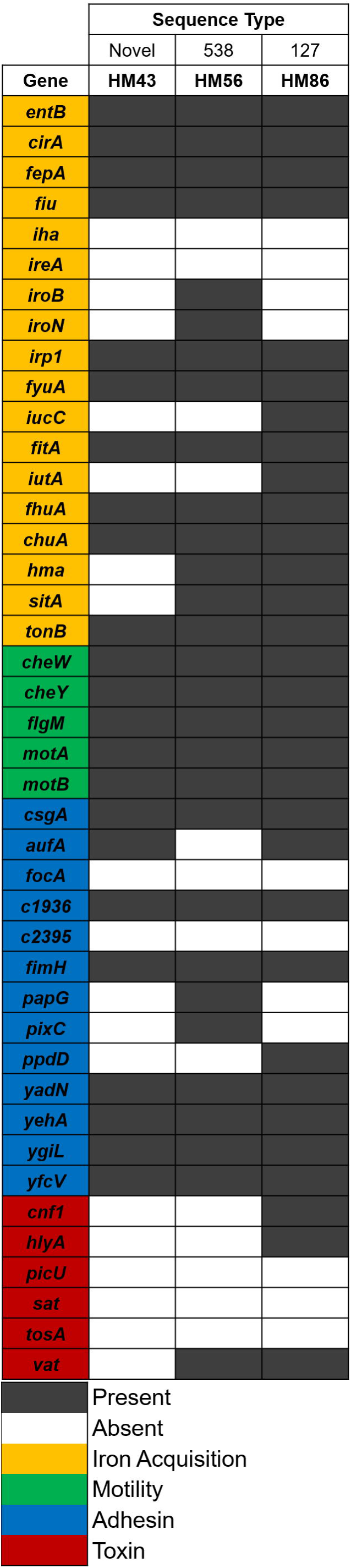
Virulence factors present in select clinical UPEC strains. Presence or absence of these 42 genes was determined via BLAST (≥80% coverage and (≥90% identity). White indicates absence of gene, while black indicates presence. Color coding on gene names identifies the function of each virulence factor. Goldenrod is iron acquisition, green is motility, blue is adhesins, while red is toxins. Sequence types were determined computationally and reported in Sintsova et al (13).

To compare UPEC gene expression during mUTI against hUTI, 40 mice were transurethrally inoculated with each UPEC strain and mouse urine was collected directly into RNAprotect, 48 hours post inoculation, for RNA isolation and sequencing. Animals were then sacrificed and the bacterial burden of their urine, bladder and kidneys was quantified. All three strains successfully colonized the animals with bacterial burdens ranging between 5.0×10^3^ – 4.4 x10^4^ CFU/g in the bladder and 1×10^4^ – 1.2×10^6^ CFU/g in the kidneys (**Fig. 2A**), levels of colonization that are consistent with an active UTI. We also assessed levels of inflammation (on a scale from 0 to 3) in the bladders and kidneys of these infected mice, comparing them to mice that were mock-infected with PBS (**Fig. 2B, Supplemental Fig.1**). After 48 hours, infection with all of the three UPEC strains resulted in mild levels of inflammation in the bladder (median inflammation scores of 1.0, 0.25, and 0.5 for HM43, HM56 and HM86, respectively) and slightly higher levels in the kidneys (median inflammation scores of 1.25, 1.5, and 1.0 for HM43, HM56 and HM86, respectively). These similar scores indicated that the general host responses were comparable across these three different strains.

**Fig. 2.**
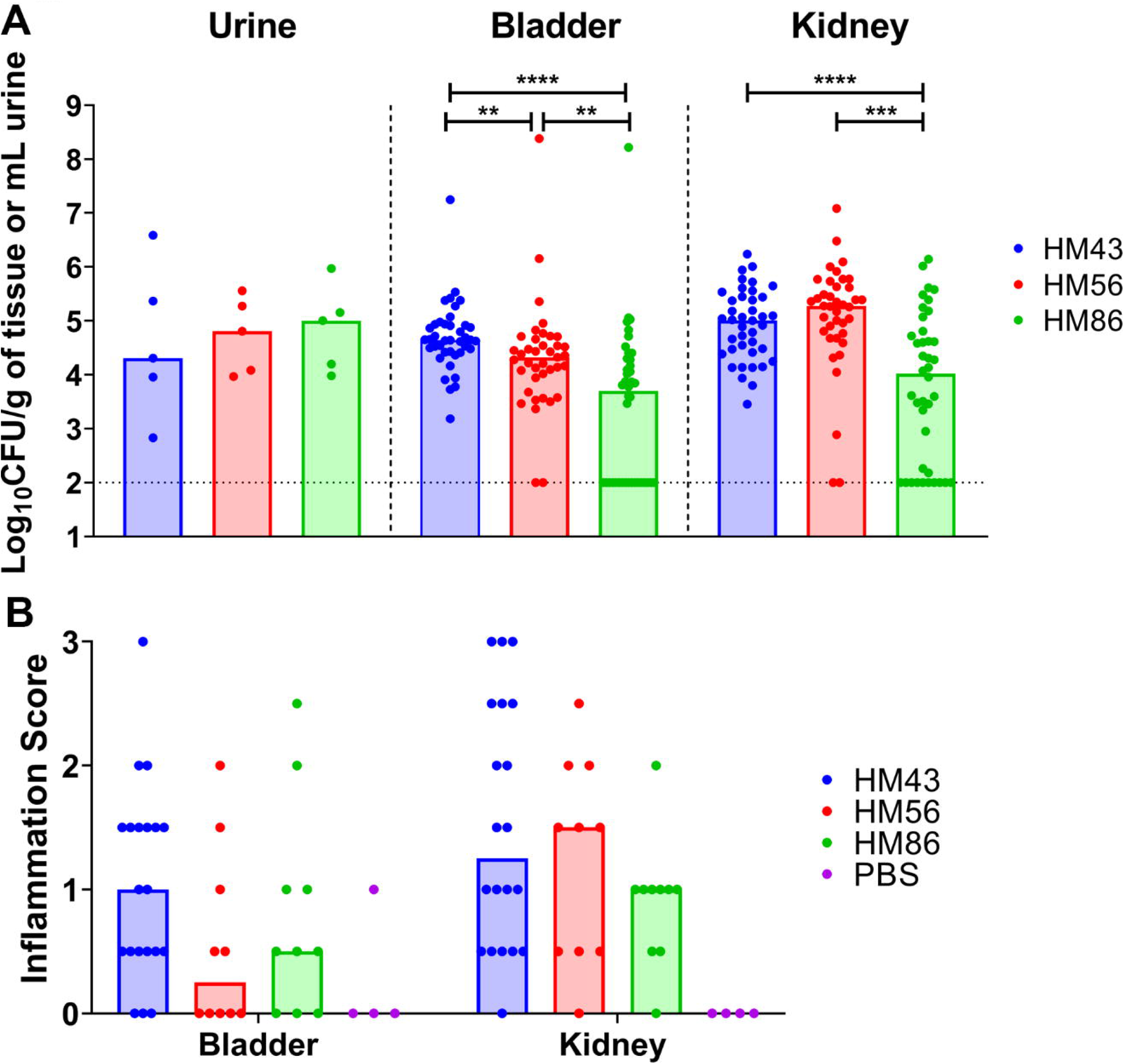
Murine colonization and inflammatory response of selected clinical UPEC strains. CBA/J mice were transurethrally inoculated with 10^8^ CFU of the indicated strain (HM43, HM56 of HM86). (A) Bacterial burden was enumerated from urine, bladder and kidneys 48 hours post infection. Symbols are individual animals and bars represent the median. Dotted line indicates limit of detection. A two-tailed Mann-Whitney test was performed to test significance, ** *P* <0.01, \*\*\**P* <0.005 \*\*\*\**P* <0.0001. (B) Inflammation was assessed using histopathological analysis of stained thin sections of each specified organ. Inflammation was scored on a 0-3 scale, with zero being no inflammation, and 3 being severe. Mice were mock-infected with PBS to serve as a negative control. Symbols are individual animals and bars represent the median.

In addition to isolating RNA from mouse urine during mUTI, we also isolated and sequenced RNA from HM43, HM56 and HM86 cultured to mid-logarithmic phase in both filter-sterilized human urine and lysogeny broth (LB). All samples processed in this study underwent identical treatments to deplete eukaryotic mRNA, prepare libraries, and conduct sequencing (see Methods).

### The bacterial transcriptome is highly correlated between human and mouse infections

First, we assessed how UPEC gene expression during hUTI compared to that during *in vitro* conditions and mUTI. For each strain, we compared log_2_ transcripts per million (TPMs) of every gene between LB and hUTI, human urine culture and hUTI, and finally mUTI and hUTI. Gene expression during hUTI and mUTI was most highly correlated with the Pearson correlation coefficient (*r*) ranging from 0.86 to 0.87 (**Fig. 3**). In contrast, the *in vitro* human urine culture when compared to hUTI exhibited lower correlation values of 0.73-0.80 (**Fig. 3)**, consistent with our previous report (13). Interestingly, gene expression correlation between LB and hUTI was higher than the correlation between urine and hUTI (*r* between 0.80-0.88) (**Fig 3**). Our data demonstrate that murine infection is the most reliable and consistent model to recapitulate the conditions that are observed during human infection.

**Fig. 3.**
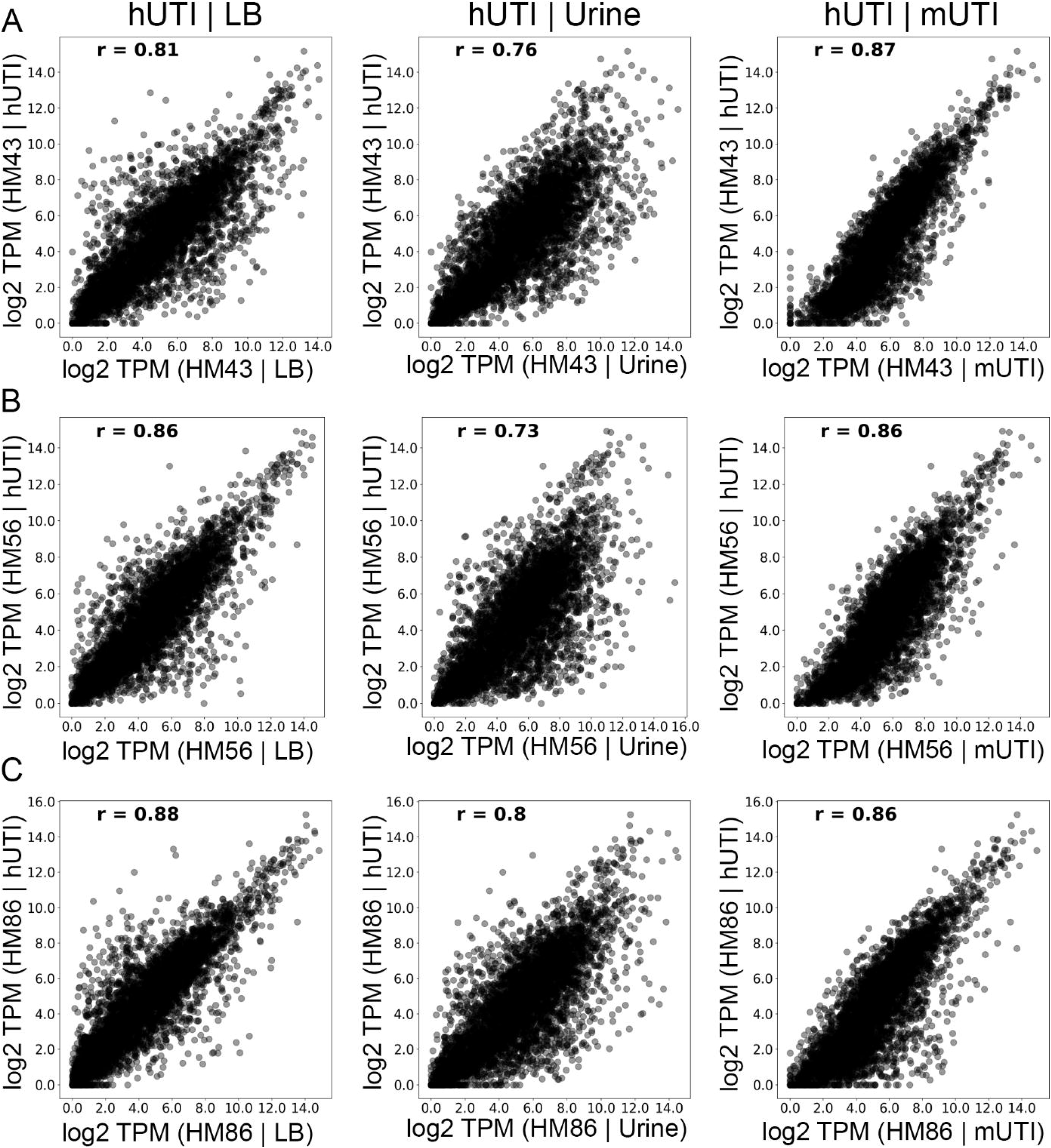
UPEC gene expression during mouse and human infections is highly correlated. Normalized gene expression (log_2_ TPM) for three UPEC strains: HM43 (A), HM56 (B), and HM86 (C) was compared between LB culture and human infection (hUTI | LB); urine culture and human infection (hUTI | urine); and mouse infection and human infection (hUTI | mUTI). Pearson correlation coefficient (*r*) is shown in top left corner of each plot.

### Contribution of growth rate to gene expression patterns

We have recently shown that diverse UPEC strains show a conserved gene expression pattern in human patients with uncomplicated UTIs (13). Since we saw such strong correlation between gene expression in patients and in mice for each of the UPEC strains (**Fig 3)**, we hypothesized that we would also observe a conserved pattern of gene expression between different UPEC strains during mUTI. To address this question, we performed principal component analysis (PCA) on gene expression of the 3,266 genes present in all three UPEC strains (**Fig. 4A**). We observed four distinct clusters that corresponded to the two *in vitro* growth conditions (LB and filter-sterilized human urine cultures) and the two infection sites (human patients and mice) all displaying condition-specific gene expression programs. Samples from patients and mice clustered closer to each other than to *in vitro* samples, suggesting that there is an infection-specific gene expression pattern conserved between the two hosts.

**Fig. 4.**
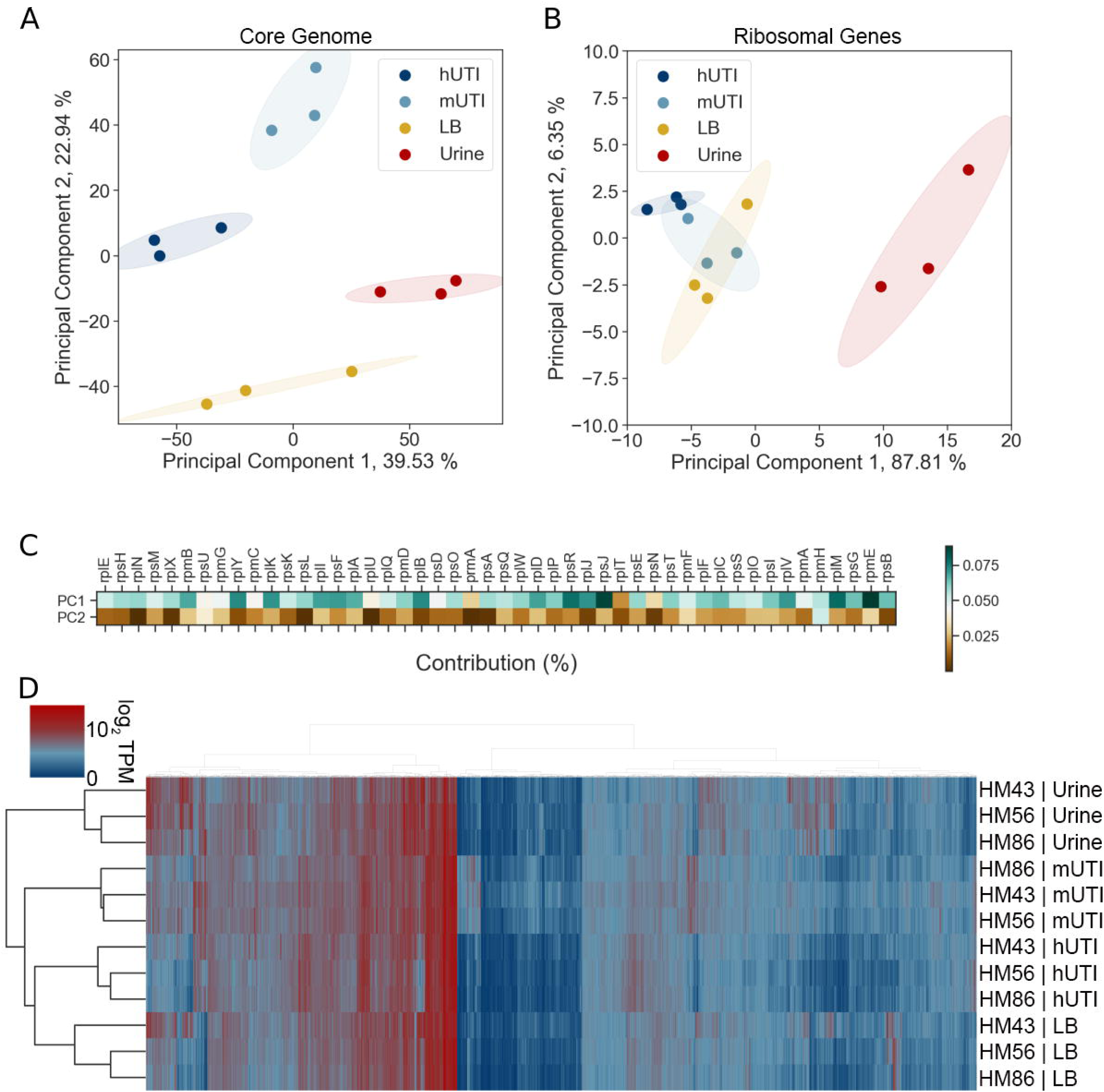
Host-associated gene expression is distinct from that of *in vitro* culture. (A) Principal component analysis of normalized gene expression of 3 clinical UPEC strains during human infection (hUTI), during mouse infection (mUTI), during *in vitro* LB culture (LB), and *in vitro* urine cultures (Urine). (B) Principal component analysis of normalized r-protein expression of 3 clinical UPEC strains during human infection (hUTI), during mouse infection (mUTI), during *in vitro* LB culture (LB), and *in vitro* urine cultures (Urine). (C) Percent contribution from r-proteins to PC1 and PC2 from PCA analysis shown in (A). (D). Hierarchical clustering of *in vitro*, murine and patient samples based on normalized gene expression of genes present in all 3 strains (n=3266).

We further wanted to estimate the contribution of growth rate to similarity of gene expression profiles between mUTI and hUTI since a major hallmark of the conserved transcriptional program of UPEC in humans is rapid growth (13-15). When cultured *in vitro*, there is no appreciable difference in the growth rate of the UPEC strains cultured in human urine (**Supplemental Figure 2A, B**). There appears to be an extremely subtle difference in growth rate in LB between HM43 and HM86 (**Supplemental Figure 2C, D**), which is interesting given that these two strains had the largest difference in correlation between hUTI and LB (*r* values of 0.81 and 0.88 respectively). However, while HM86 had the highest correlation, it grew more slowly than HM43, indicating that it is potentially more than just growth rate that is driving this similarity.

We also performed PCA using only ribosomal protein expression data (**Fig. 4B**), since expression of ribosomal proteins is directly correlated with bacterial growth rate (16, 17) as another method to dissect the contribution of growth rate. While this analysis still showed a clear difference between urine samples and other conditions, it failed to clearly separate LB, mUTI and hUTI samples (**Fig. 4B**). This suggests that while growth rate undoubtedly shapes the gene expression pattern in mUTI and hUTI, growth rate alone is insufficient to explain gene expression patterns observed in **Fig. 4A** and **Fig. 3**. Additionally, we looked at how much each of the ribosomal genes contributed to the either Principal Component 1 (PC1) or Principal Component 1 (PC2) (**Fig. 4C**). We found that all of the ribosomal genes contributed more to PC1 than PC2 values. Thus, we hypothesize that PC1 might in fact separate samples based on growth rate, while other unknown factors account for sample separation along PC2.

The conclusions of PCA analysis (clear separation of samples based on condition rather than on strain) were independently confirmed using Ward’s hierarchical clustering of log_2_ TPM values (**Fig 4D**). We also demonstrate that growth in rich medium (LB) more closely mimics hUTI than growth in nutrient-poor human urine, in agreement with the previously demonstrated rapid growth of UPEC in the host (both mouse and human) compared to slow growth in human urine (13-15, 18).

### Differentially regulated genes between human and mouse infection suggest nutritional disparities

Despite the high concordance of hUTI and mUTI gene expression data, we wanted to determine whether any genes are differentially regulated between human and mouse infection. To answer this question, we used the R package DEseq2 (19) to find significant differences in gene expression between the two different infections. Strikingly, only 175 genes, representing 5.4% of the 3,266 genes analyzed, were differentially regulated (30 upregulated, 145 downregulated) in human infection compared to the mouse model (**Fig. 5A, Table 1** and **2**, **Supplemental Table 2**).

**Table 1:** Genes differentially upregulated in human patients compared to mice^a^.

| Gene | Annotation | log<sub>2</sub> FC | Locus Tag |
| --- | --- | --- | --- |
| ***ahpC*** | alkyl hydroperoxide reductase, AhpC component | 2.6 | b0605 |
| ***cspA*** | cold shock protein CspA | 4.3 | b3556 |
| ***dsdX*** | D-serine transporter | 2.7 | b2365 |
| ***fis*** | DNA-binding transcriptional dual regulator Fis | 2.4 | b3261 |
| ***ftsB*** | cell division protein FtsB | 2.3 | b2748 |
| ***gntK*** | D-gluconate kinase, thermostable | 2.7 | b3437 |
| ***gpt*** | xanthine-guanine phosphoribosyltransferase | 2.4 | b0238 |
| ***gspH*** | hypothetical type II secretion protein GspH | 3.0 | UTI89_C3381 |
| ***gspL*** | hypothetical type II secretion protein GspL | 3.3 | UTI89_C3377 |
| ***hpt*** | hypoxanthine phosphoribosyltransferase | 2.1 | b0125 |
| ***ibaG*** | acid stress protein IbaG | 2.2 | b3190 |
| ***lysP*** | lysine:H(+) symporter | 2.1 | b2156 |
| ***opgC*** | protein required for succinyl modification of osmoregulated periplasmic glucans | 2.6 | b1047 |
| ***ribE*** | 6,7-dimethyl-8-ribityllumazine synthase | 2.1 | b0415 |
| ***rpmE*** | 50S ribosomal subunit protein L31 | 2.7 | b3936 |
| ***suhB*** | inositol-phosphate phosphatase | 2.6 | b2533 |
| ***yajG*** | putative lipoprotein YajG | 2.6 | b0434 |
| ***yceA*** | UPF0176 protein YceA | 3.5 | b1055 |
| ***yciB*** | inner membrane protein | 2.3 | b1254 |
| ***ydiE*** | PF10636 family protein YdiE | 2.1 | b1705 |
| ***yecJ*** | DUF2766 domain-containing protein YecJ | 2.5 | b4537 |
| ***yejL*** | DUF1414 domain-containing protein YejL | 2.2 | b2187 |
| ***yfaZ*** | putative porin YfaZ | 2.9 | b2250 |
| ***yfhL*** | putative 4Fe-4S cluster-containing protein YfhL | 2.6 | b2562 |
| ***yghD*** | putative type II secretion system M-type protein | 3.6 | b2968 |
| ***yghG*** | lipoprotein YghG | 3.3 | b2971 |
| ***yifK*** | putative transporter YifK | 3.6 | b3795 |
| ***yqcC*** | DUF446 domain-containing protein YqcC | 2.5 | b2792 |
| ***yqgF*** | ribonuclease H-like domain containing nuclease | 2.3 | b2949 |
| ***yqiA*** | esterase YqiA | 2.3 | b3031 |

<sup>a</sup> FC: Fold Change

**Table 2:** Top 30 genes differentially downregulated in human patients compared to mice^a^.

| Gene | Annotation | log<sub>2</sub> FC | Locus Tag |
| --- | --- | --- | --- |
| *allB* | allantoinase | -7.0 | b0512 |
| *allD* | ureidoglycolate dehydrogenase | -6.0 | b0517 |
| *araA* | L-arabinose isomerase | -5.6 | b0062 |
| *citC* | citrate lyase synthetase | -6.2 | b0618 |
| *citD* | citrate lyase acyl carrier protein | -5.6 | b0617 |
| *citG* | triphosphoribosyl-dephospho-CoA synthase | -5.5 | b0613 |
| *citX* | apo-citrate lyase phosphoribosyl-dephospho-CoA transferase | -5.9 | b0614 |
| *eutG* | putative alcohol dehydrogenase in ethanolamine utilization | -5.7 | b2453 |
| *eutM* | putative structural protein, ethanolamine utilization microcompartment | -6.1 | b2457 |
| *eutN* | putative carboxysome structural protein | -6.6 | b2456 |
| *fdrA* | putative acyl-CoA synthetase FdrA | -6.2 | b0518 |
| *frdA* | fumarate reductase flavoprotein subunit | -5.6 | b4154 |
| *frdB* | fumarate reductase iron-sulfur protein | -5.3 | b4153 |
| *frdC* | fumarate reductase membrane protein FrdC | -6.1 | b4152 |
| *glxR* | tartronate semialdehyde reductase 2 | -7.7 | b0509 |
| *hycA* | regulator of the transcriptional regulator FhlA | -5.3 | b2725 |
| *hycB* | formate hydrogenlyase subunit HycB | -6.0 | b2724 |
| *ompW* | outer membrane protein W | -6.4 | b1256 |
| *ssnA* | putative aminohydrolase | -5.3 | b2879 |
| *tdcA* | DNA-binding transcriptional activator TdcA | -6.2 | b3118 |
| *tdcB* | catabolic threonine dehydratase | -5.5 | b3117 |
| *ulaA* | L-ascorbate specific PTS enzyme IIC component | -6.1 | b4193 |
| *ulaB* | L-ascorbate specific PTS enzyme IIB component | -5.8 | b4194 |
| *ulaC* | L-ascorbate specific PTS enzyme IIA component | -5.4 | b4195 |
| *ybbW* | putative allantoin transporter | -6.7 | b0511 |
| *ygeW* | putative carbamoyltransferase YgeW | -6.9 | b2870 |
| *ygeY* | putative peptidase YgeY | -6.1 | b2872 |
| *ygfK* | putative oxidoreductase, Fe-S subunit | -5.8 | b2878 |
| *yhjX* | putative pyruvate transporter | -5.4 | b3547 |
| *ylbE* | DUF1116 domain-containing protein YlbE | -5.6 | b4572 |

<sup>a</sup> FC: Fold Change, N/A: not available

**Figure 5.**
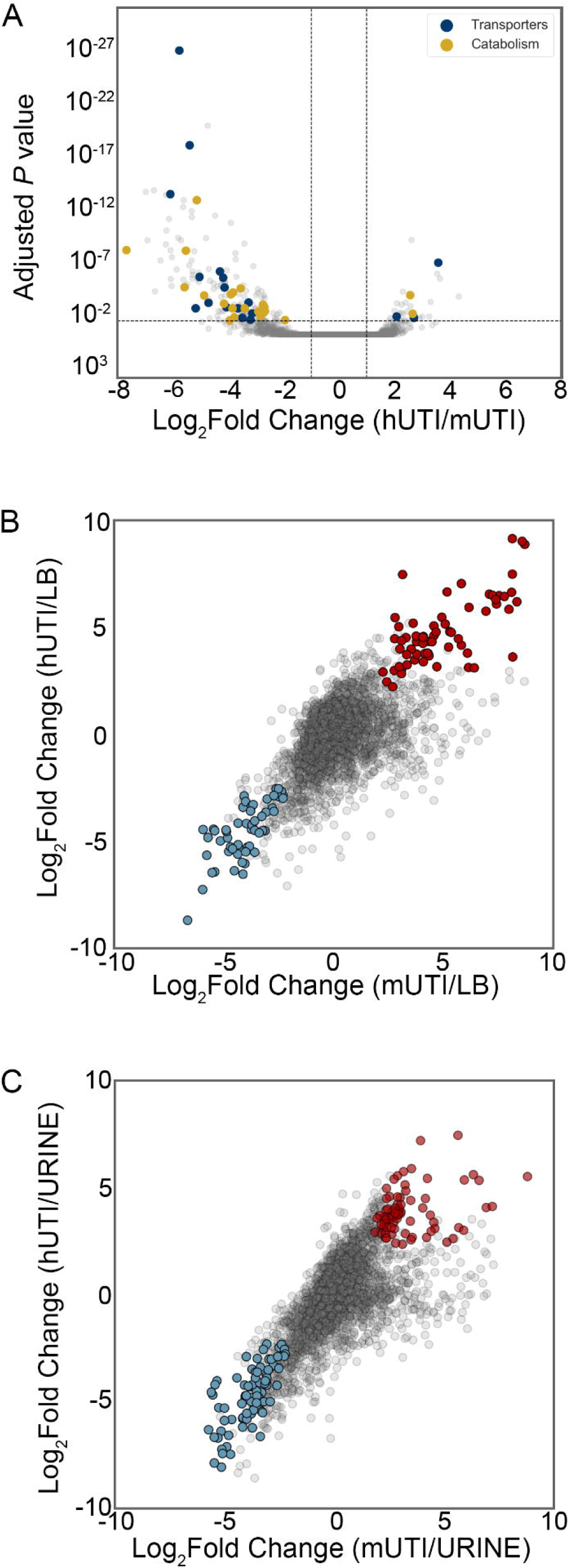
Differential expression analysis reveals infection-specific gene expression responses. (A) The DESeq2 R package was used to compare UPEC gene expression during m UTI to that in patients. Each UPEC strain was considered an independent replicate (n = 3). Genes were considered up-regulated (down-regulated) if log_2_ fold change in expression was higher (lower) than 1 (vertical lines), and *P* value < 0.05 (horizontal line). Using these cutoffs, we identified 30 upregulated genes and 145 downregulated genes in patients. GO/pathway analysis showed a number of transporters and catabolic enzymes among differentially expressed genes. (B and C) Identification of genes differentially expressed during infection (hUTI or mUTI) compared to LB (B) or urine (C). Genes were considered to be up/downregulated in both mouse and human if log_2_ fold change was higher/lower than 1, and *P* value < 0.05 in both cases. Genes that were upregulated during infection when compared to LB (B) or urine (C) are shown in red, genes that were downregulated during infection compared to LB are shown in blue.

The upregulated gene with the highest log_2_ fold-change difference (4.3) between human and mouse was *cspA*, which encodes an RNA chaperone initially identified as a cold shock protein (20, 21). However, this protein may have other functions as it is highly expressed during early exponential phase (22) and during the introduction of fresh nutrient sources (23). In addition, one of the genes responsible for cell division, *ftsB*, and a major regulator of ribosomal RNA transcription, *fis*, were upregulated in hUTI as compared to mUTI.

The majority of differentially regulated genes were downregulated in patients compared to mice, and several of these genes span operons encompassing specific systems (**Table 2** and **Supplemental Table 2**). For example, during mUTI, we observed: increased expression of the citrate lyase operon *citCDEFGTX*, which is responsible for the conversion of citrate oxaloacetate and acetate and feeds into the production of acetyl-COA under anaerobic conditions (24); the pathway for allantoin breakdown (*allABDC*); and ethanolamine utilization (*eutABCDEGHJLMNPQST*) (**Table 2** and **Supplemental Table 2**). In addition, genes encoding transporters for the uptake of L-arabinose (*araADFGH)*, L-ascorbate (*ulaABCDEDF*), and allantoin (*ybbW*) were transcribed at higher levels during mUTI (**Table 2** and **Supplemental Table 2**). Furthermore, several genes related to anaerobic metabolism or fermentation (*hycBDF* and *frdAB)* were more highly expressed in mice (**Table 2** and **Supplemental Table 2**). All of these results indicate subtle nutrient differences between the mouse and human urinary tract.

### Infection-specific gene expression

Additionally, we were interested in identifying genes that behaved similarly during both human and mouse infection, *i.e*., genes that were up- or down-regulated in both mouse and human UTI when compared to either of the *in vitro* conditions (LB or filter-sterilized human urine). There were 54 downregulated genes in both mouse and human UTI when compared to LB (**Fig. 5B, Supplemental Table 3**) and there were 67 upregulated genes during both mUTI and hUTI when compared to LB (**Fig. 5B, Supplemental Table 4**). Interestingly, both chemotaxis (*cheABWYZ*) and flagellar machinery *(flgCFGLM* and *fliS*) were downregulated during infection, which may be attributed to the fact that the UPEC strains we are analyzing were isolated from the urine of infected individuals; motility genes tend to be upregulated when UPEC enters the ureters to ascend to the kidneys (25). In contrast, *nrdEFHI* genes are upregulated in both mice and humans compared to LB. These genes are ribonucleotide reductases required for DNA synthesis, and therefore often associated with fast growth, fitting the previously established paradigm of UPEC’s rapid *in vivo* growth rate during human and murine infection (13-15, 18).

There were 82 genes that were downregulated during either mUTI or hUTI when compared to urine (**Fig. 5C, Supplemental Table 5**). These included branched-chain amino acid biosynthesis (*ilvCDEMN*) and leucine biosynthesis (*leuABCD*) operons, consistent with previous literature indicating that UPEC scavenges amino acids and peptides during infection(13, 26). In contrast, there were 72 genes that were upregulated in humans and mice when compared to urine (**Fig. 5C, Supplemental Table 6**). As previously reported (13), we observed 16 genes associated with ribosomal subunit production as well as the master regulator *fis*, which activates rRNA transcription, together reinforcing our observation that bacteria are growing rapidly in the host (13, 14).

### Expression of fitness factors during murine infection is predictive of gene expression during human infection

Finally, we wanted to determine whether previously identified UPEC virulence factors that have been studied using *in vivo* mouse models would show comparable levels of expression during both mouse and human infections. We focused on three major functional groups of fitness factors: iron acquisition systems, adhesins, and metabolism (**Supplemental Table 7**). We plotted log_2_ TPM of the genes in each functional group, comparing expression between hUTI and LB, hUTI and urine, as well as hUTI and mUTI for each of the UPEC strains (**Fig. 6, Supplemental Fig. 3, Supplemental Fig 4**.)

**Figure 6.**
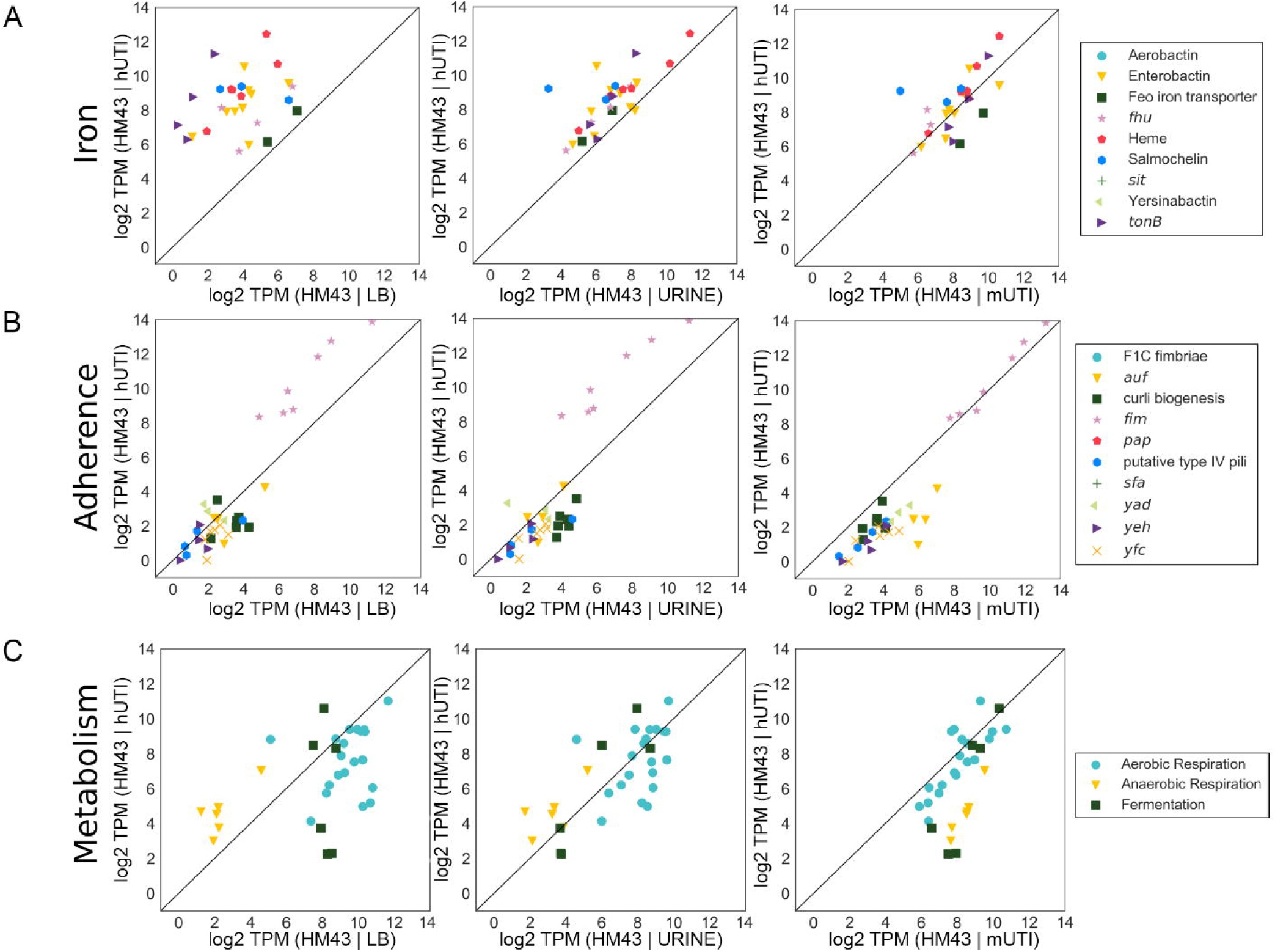
Gene expression of virulence factors as well as metabolic machinery is highly consistent between mouse model of UTI and human UTI. Normalized gene expression of iron acquisition operons (A), adherence genes (B), and metabolic pathways (C) for HM43 was compared between LB and human infection (LB | hUTI), urine and human infection (urine | hUTI), and mouse UTI and human UTI (mUTI | hUTI).

As expected, iron acquisition gene expression is much higher during human infection than during growth in rich LB medium (**Fig. 6A**). The expression levels are more similar between urine culture (an iron-poor medium) and human infection, but murine infection provides the most analogous profile (**Fig. 6A**). The only adherence gene cluster that was highly expressed in any of the assayed conditions was the *fim* operon, which encodes type 1 fimbriae. Expression of *fim* genes was higher in patients compared to either of the *in vitro* conditions, but almost perfectly matched the expression levels observed during mouse infection (**Fig. 6B**).

Anaerobic metabolic genes showed a major difference between human infection and *in vitro* growth. The converse is also true; aerobic respiration genes, in particular, were expressed at lower levels in humans than in either *in vitro* condition. Importantly, we also observed that the expression levels of aerobic respiration genes align concordantly between human and mice (**Fig. 6C**), while anaerobic respiration gene expression was elevated in mice compared to humans. This observation corroborates results from **Fig. 5A, Table 2** and **Supplemental Table 2**, where several genes involved in anaerobic metabolism were expressed at higher levels in mice compared to hUTI. Overall, we conclude, with only limited exceptions, that the mouse model of UTI not only shows a strong global correlation of gene expression with hUTI, but also closely reflects the expression of virulence and fitness genes that are known to contribute to UPEC fitness during human infection.

## Discussion

UPEC virulence factors as well as potential therapeutic strategies have been studied in detail using a well-established mouse model of ascending UTI that involves transurethral inoculation of UPEC into the bladder. This mouse model has been extensively used in the field, and the original papers defining this model (8, 27) have been cited nearly 500 times. While this model has been the gold-standard in the field to relate scientific discovery to human health, there are some differences between the experimental model and what occurs in human infection. In the murine model >10^7^ CFU are inoculated directly into the mouse bladder, while in human infection the number of infecting bacteria is likely to be far lower, and bacteria are not directly introduced into the bladder. Rather, in humans, it is thought that the periurethral region is transiently colonized by UPEC, and bacteria ascend to the bladder and in some cases to the kidneys (28). Furthermore, there are some differences in the immune responses between mice and humans. For example, mice express Toll-like receptor 11 (TLR11), which induces host inflammation in response to UPEC and provides the kidneys with a modest level of protection from bacterial colonization (29). However, humans do not express TLR11 (29), indicating that not all murine responses will precisely mimic the human response. Therefore, it is of the utmost importance to validate the murine model of UTI and understand the extent to which it recapitulates human disease. Until now, there has been no direct comparison of global bacterial gene expression between human and mouse studies using the identical strain, and it is essential to understand how the mouse model relates to human disease. This study is the first to define the bacterial transcriptome from infected patients and infected mice using the same UPEC strain, thus presenting a direct comparison between the murine model and human infection. Our study demonstrates that the UTI mouse model accurately recapitulates the human disease with respect to the bacterial transcriptional response.

We compared three UPEC strains (HM43, HM56, and HM86) that were isolated in 2012 from women with symptoms of cystitis and documented significant bacteriuria (30). We isolated bacterial RNA, stabilized immediately, from their urine to conduct RNA-seq and define the core bacterial transcriptome during acute human infection (13). The same strains were then used for mouse infection, followed by urine collection, bacterial RNA isolation and sequencing. We consistently observed an extraordinarily high correlation between the bacterial transcriptome during mouse and human infection, with the Pearson correlation coefficient ranging from 0.86 to 0.87. This correlation is not strain-specific, as infections with all three stains showed similar results. Expression of virulence and metabolic genes was also found to be very similar between human and mouse infection. While all three strains belonged to B2 phylogroup, we do not believe this to be a phylogroup-specific effect as we have previously shown few differences in gene expression between different UPEC phylogroups (13). This provides strong evidence that the mouse model is an accurate representation of the infection that occurs in humans, and by understanding which genes do not correlate between human and mouse, we can further understand the limitations of the mouse model.

Mounting evidence suggests that UPEC in the human host are rapidly dividing (13, 14), and we have recently shown that this rapid growth rate is recapitulated in the mouse model, although to a lesser degree (13). This difference in growth rate between human and mouse UTIs potentially can be understood by examining the genes that are differentially expressed between human and mouse infections. Most of the differentially expressed genes were expressed at a lower level during hUTI compared to mUTI (145 of 175 genes). Many of these 145 genes are involved in anaerobic metabolism. Several of them were clustered in operons encoding oxidoreductases involved in fumarate or nitrite reduction. Additionally, genes involved in nutrient usage under anaerobic conditions were also expressed at a lower level during hUTI, such as the *all* operon, which encodes the catabolic pathway for allantoin degradation (a step in purine catabolism (31)), or the *ula* operon, which encodes both an L-ascorbate transporter and the corresponding enzymes for L-ascorbate utilization, a compound that can be present in urine due to its water soluble nature (32). Therefore, we hypothesize that the human bladder is better oxygenated than the mouse bladder. Indeed, a higher oxygen level in human bladders might also account for the higher levels of replication observed in hUTI (13), since an aerobic lifestyle can support more rapid growth. There were also differences in transport systems involved in nutrient acquisition between hUTI and mUTI. Arabinose transport (*araADFGH*) was expressed at a lower level in humans; this sugar has been shown to be present in human bladders in μM amounts when normalized to creatine (32). It is also present in murine bladders (33), but has never been precisely quantified. It would be interesting if arabinose is present in lower amounts in humans compared to mice, accounting for this difference in regulation. One of the few genes that was upregulated in humans compared to mice was *dsdX*, which encodes a D-serine transporter. Interestingly, D-serine is present in micromolar amounts in human urine (34), D-serine utilization is associated with uropathogenic strains (35, 36) and accumulation of D-serine leads to a “hypervirulent” phenotype (36, 37) in the urinary tract. Since there was an upregulation in D-serine transport, and not in the deaminase required for its breakdown (*dsdA*), perhaps this presents a mechanism to increase the intracellular levels of D-serine specific to hUTI.

We also compared the gene expression profiles of UPEC during infection with the two most common *in vitro* models, LB cultures and pooled filter-sterilized human urine cultures. Surprisingly, even though urine might seem to be the more physiologically relevant medium for *in vitro* experimentation, LB overall provided a better model for infection compared to urine cultures. Indeed, the correlations of strains grown in LB approached the correlations when comparing human infection to the mouse model (**Fig. 3**). However, pathway analysis using the online tool DAVID (38) revealed several statistically significantly enriched (enrichment score ≥1.5) pathways that were enriched when comparing either hUTI with LB (**Supplemental Table 8**) or mUTI (**Supplemental Table 9**). Several pathways were shared in both comparisons, specifically iron homeostasis or acquisition, as well as motility and various metabolism pathways (TCA, arginine and proline, and sulfur) were differentially regulated between the LB and *in vivo* conditions. (**Supplemental Table 8** and **9**). Not only do these results corroborate the results from **Fig. 6** they also point to shared differences when comparing either infection model to *in vitro* further underscoring the similarities between mUTI and hUTI.

A major difference between *in vitro* growth of UPEC in LB or urine is growth rate, and one of the major hallmarks of the transcriptional program of UPEC during infection is rapid growth. Subsequently, when comparing the pathways enriched in the differentially regulated genes between hUTI or mUTI to urine, ribosomes are enriched (**Supplemental Table 10** and **11**). However, that is just one of seven other statistically significantly pathways that are enriched; there is a variety of factors influencing this outcome. Furthermore, several of these pathways (biosynthesis of amino acids, flavin adenine dinucleotide binding, and nitrogen metabolism/nitrogen utilization) are shared between mUTI vs urine and hUTI vs urine, implying that there are similarities between the two infections that are not observed in *in vitro* growth in urine (**Supplemental Table 10** and **11**).

There are several factors that vary between infection conditions and *in vitro* growth that could account for the lower correlation between hUTI and urine culture gene expression. For example, a nutrient limitation specific to *in vitro* growth. The bladder is akin to a chemostat, with fresh urine constantly being introduced into the organ, a condition that is not recapitulated in the *in vitro* conditions. Furthermore, the collected urine is typically filter-sterilized and this method excludes exfoliated bladder epithelial cells, likely another major source of nutrients for the pathogen during human infection. In future studies, we could add lysed bladder cells from cell culture to supplement the filter-sterilized urine and determine if this represents a better model of the nutrient milieu. However, when studying specific systems, such as iron acquisition, urine is a better model than LB, since it more accurately recapitulates the iron-limited environment of the host.

In summary, while both *in vitro* models have advantages and disadvantages, the mouse model offers a holistic representation of infection and accurately models UPEC pathway expression. Detailed examination of gene expression profiles between mice and humans allows us to understand which parts of the model are not recapitulated in patients, so we may tailor our studies accordingly. This way the mouse model can serve as a platform to answer questions that are more difficult or impossible to assess when working with human patients.

## Methods

### Bacterial culture conditions

Clinical UPEC strains HM43, 56 and 86 (30) were cultured overnight in LB medium at 37°C with aeration. The next morning, cultures were centrifuged and the pellets washed twice with PBS, then diluted 1:100 into either fresh LB medium or human urine. The human urine was collected and pooled from at least four healthy female volunteers and passed through a 0.22 μm filter for sterilization. Bacteria were cultured at 37°C with aeration to mid-exponential phase (∼3 hours), then stabilized in RNAprotect (Qiagen). Bacterial pellets were stored at −80°C until RNA isolation.

### Mouse infection

Forty female CBA/J mice were transurethrally inoculated, using the previously established ascending model of UTI (8), with 10^8^ CFU of either HM43, 56, or 86 that had been cultured in LB overnight with shaking. The infection was allowed to progress for 48 hours. Urine from five mice was collected to enumerate bacterial burden, while the rest was collected for RNA (see below for method). Mice were sacrificed, and their bladders and kidneys aseptically removed, homogenized, and plated to determine bacterial burden. Mouse urine was collected as previously described (13). Briefly, 48 HPI, urine was directly collected into RNAprotect, pooled, and pelleted. This was repeated every 45 minutes five more times resulting in a total of six pellets. These pellets were stored at −80°C until RNA isolation.

### RNA isolation and library preparation

RNA was isolated as previously described (13). Briefly, all bacterial pellets were treated with both lysozyme and proteinase K, and then total RNA was extracted using a RNeasy kit (Qiagen). Genomic DNA was removed using the Turbo DNA-free kit (ThermoFisher). Eukaryotic mRNA was depleted using Dynabeads covalently linked with oligo dT (ThermoFisher). The *in vitro* samples underwent the same treatment with Dynabeads to reduce any potential biases this procedure might introduce to the downstream sequencing. The supernatant was collected from this treatment, and RNA was concentrated and re-purified using the RNA Clean and Concentrator kit (Zymo).

To compare the results of the new RNA-sequencing experiment to the published expression data obtained from the human samples (13), library preparation method needs to be identical to avoid batch effects. The original sequencing data (hUTI) were obtained using the ScriptSeq Complete kit (Bacteria) to prepare the cDNA library. However, at the time of this study, Illumina had discontinued this kit. Therefore, we needed to use any leftover kits that had the same base preparation method (Scriptseq). To accomplish this, we used ScriptSeq Complete Gold Kit (Epidemiology), which also contains rRNA removal for bacteria and eukaryotes for the HM86 mouse sample, and all three HM43 samples (mouse, LB, and urine). ScriptSeq Complete (Bacteria) was used on the HM56 mouse sample, where the kit contained rRNA removal for bacteria; mammalian rRNA was removed with ThermoFisher’s mammalian rRNA removal kit (cat #457012). Then the *in vitro* samples from both HM56 and HM86 were prepared using the ScriptSeq Complete (bacteria) kit, which removes bacterial rRNA.

### RNA-sequencing

*E. coli* HM43 was sequenced using an Illumina HiSeq2500 (single end, 50 bp read length) and *E. coli* HM56 and HM86 were sequenced using the Nextseq-500 with identical conditions (single end, 50 bp read length).

### Histology and tissue processing

Bladders were removed and halved by cutting on the transverse plane, while kidneys were cut on the sagittal plane. One half of each organ was used to enumerate CFU, while the other halves were placed into tissue cassettes and immersion-fixed in 10% formalin for at least 24 hours. They were then embedded in paraffin, cut into thin sections and stained with hematoxylin and eosin (H&E) by the *In Vivo* Animal Core at the University of Michigan. Tissue sections were scored as described in **Table 3**. Briefly, this is a scale of 0-3, where 0 was no inflammation, and 3 was severe inflammation. Each organ section was scored by two different people in a blinded manner and the scores averaged together.

**Table 3:** Scoring criteria for histopathological analysis.

| **Table 3: Scoring criteria for histopathological analysis.** |  |  |  |  |
| --- | --- | --- | --- | --- |
|  | **Score** |  |  |  |
| **Lesion** | **0** | **1** | **2** | **3** |
| Cystitis | No scorable lesions | very rare PMNs in stroma or lumen or occasional perivascular lymphoid cuffs | many PMNs and moderate edema | Many PMNs; widespread, marked edema, transmural inflammation |
| Pyelonephritis | No scorable lesions | very occasional PMNs in lumen or peripelvic tissue | Rafts of PMNs in the pelvis and/or scattered focal aggregates of PMNs in peripelvic tissue | many PMNs in all sections, or a single large focus of PMNs in one section |

### RNAseq Data Processing

A custom bioinformatics pipeline was used for the analysis (github.com/ASintsova/rnaseq_analysis). Raw fastq files were processed with Trimmomatic (21) to remove adapter sequences and analyzed with FastQC to assess sequencing quality. Mapping was done with bowtie2 aligner (39) using default parameters. Alignment details can be found in **Supplemental Table 1**. Read counts were calculated using HTseq htseq-count (40).

### Pearson correlation coefficient calculation, PCA analysis and hierarchical clustering analysis

For PCA and correlation analysis, transcript per million (TPM) was calculated for each gene; TPM distribution was then normalized using log_2_ transformation and these normalized data were used for both PCA and correlation, and hierarchical clustering analysis. Pearson correlation and PCA were performed using Python sklearn library. Hierarchical clustering was performed using Ward’s method and Euclidean distance. All of the analysis we also performed using centered log-ratio transformation (instead of log_2_ transformation), which has recently proposed for compositional data and saw similar results (these results can be found in Jupyter notebook associated with this manuscript). Jupyter notebooks used to generate the figures are available at https://github.com/ASintsova/upec_mouse_model.

### Differential expression analysis

Differential expression analysis was performed using DESeq2 R package (19). Genes with log_2_ fold change of greater than 1 or less than −1 and adjusted *p* values (Benjamini-Hochberg adjustment) of less than 0.05 were considered to be differentially expressed. Pathway analysis was performed using R package topGO (41).

### Data access

Jupyter notebooks as well as all the data used to generate the figures in this paper are available on github: https://github.com/ASintsova/upec_mouse_model.

