## Supplemental Fig 1-4 for "The Gene Expression Profile of Uropathogenic *Escherichia coli* in Women with Uncomplicated Urinary Tract Infections Is Recapitulated in the Mouse Model"

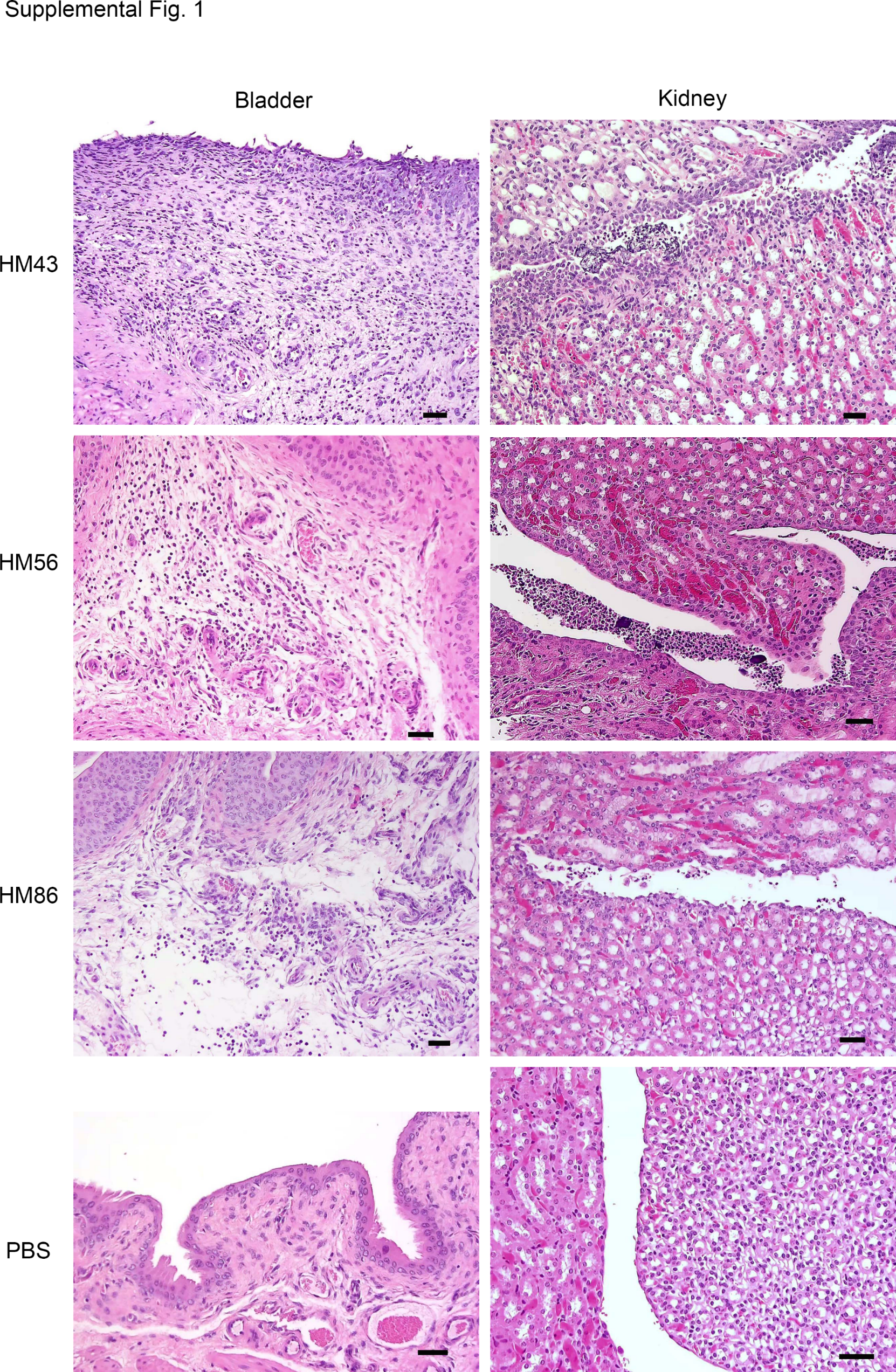

**Supplemental Figure 1. H&E stained thin sections of bladders and kidneys of infected mice.** Mice were infected with indicated UPEC strain (HM43, HM56 or HM86), or mock-infected with PBS. Bladder or kidney tissue was thin-sectioned and stained with H&E. Scale bar indicates 40 μm.

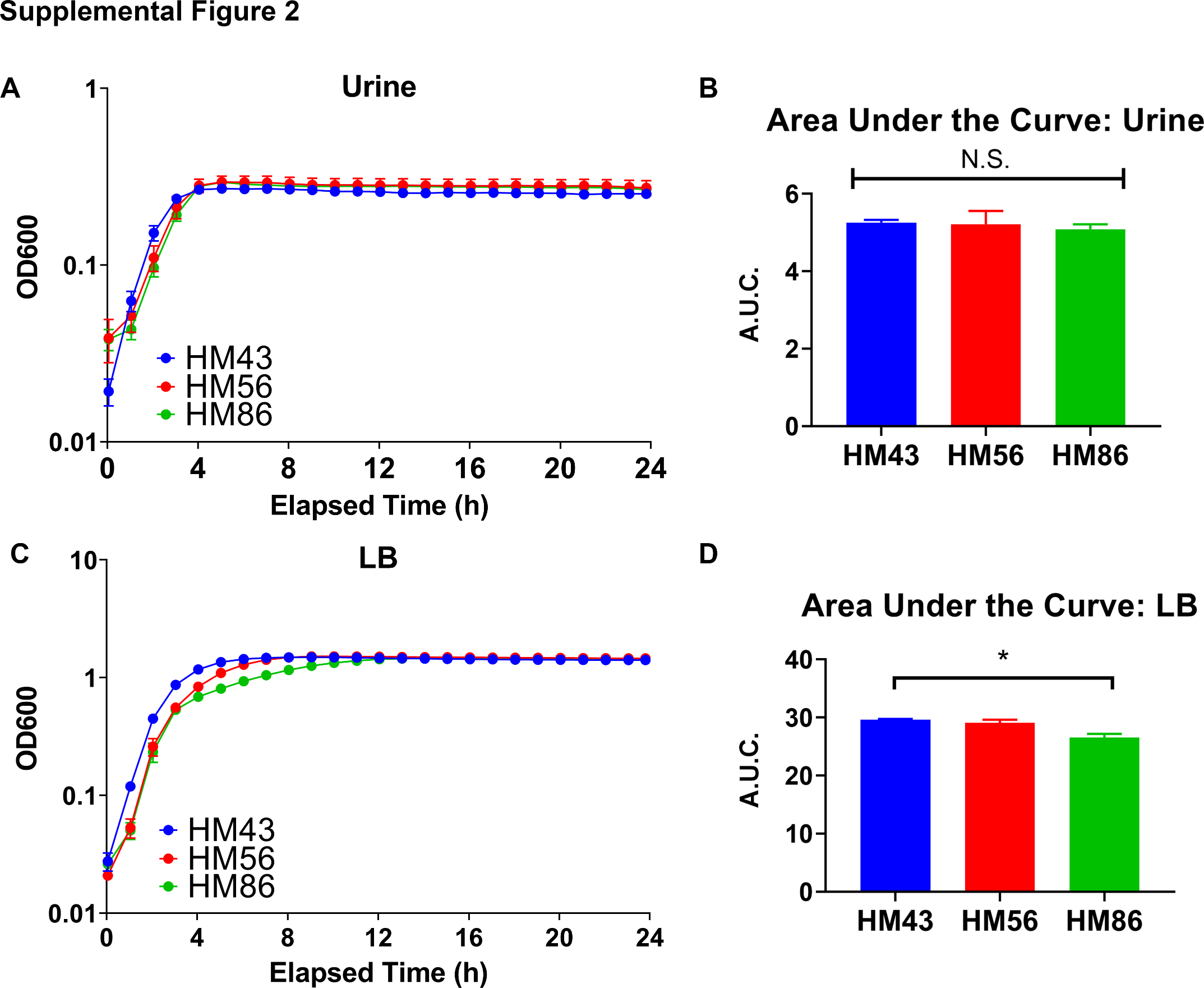

**Supplemental Figure 2. *In vitro* growth curves of strains HM43, 56 and 86.** Each strain was either grown in pooled human urine **(A)** or LB **(C)** for 24 hours. Each curve represents four independent replicates; error bars are ±Standard Error of Mean (SEM). (**B**) and (**D**) are the area under the curve (A.U.C.) of each strain from either urine or LB, respectively. A.U.C was calculated in GraphPad Prism. A.U.C. values were compared with nonparametric one-way ANOVA with Dunn’s correction for multiple testing. *, *P*<0.05

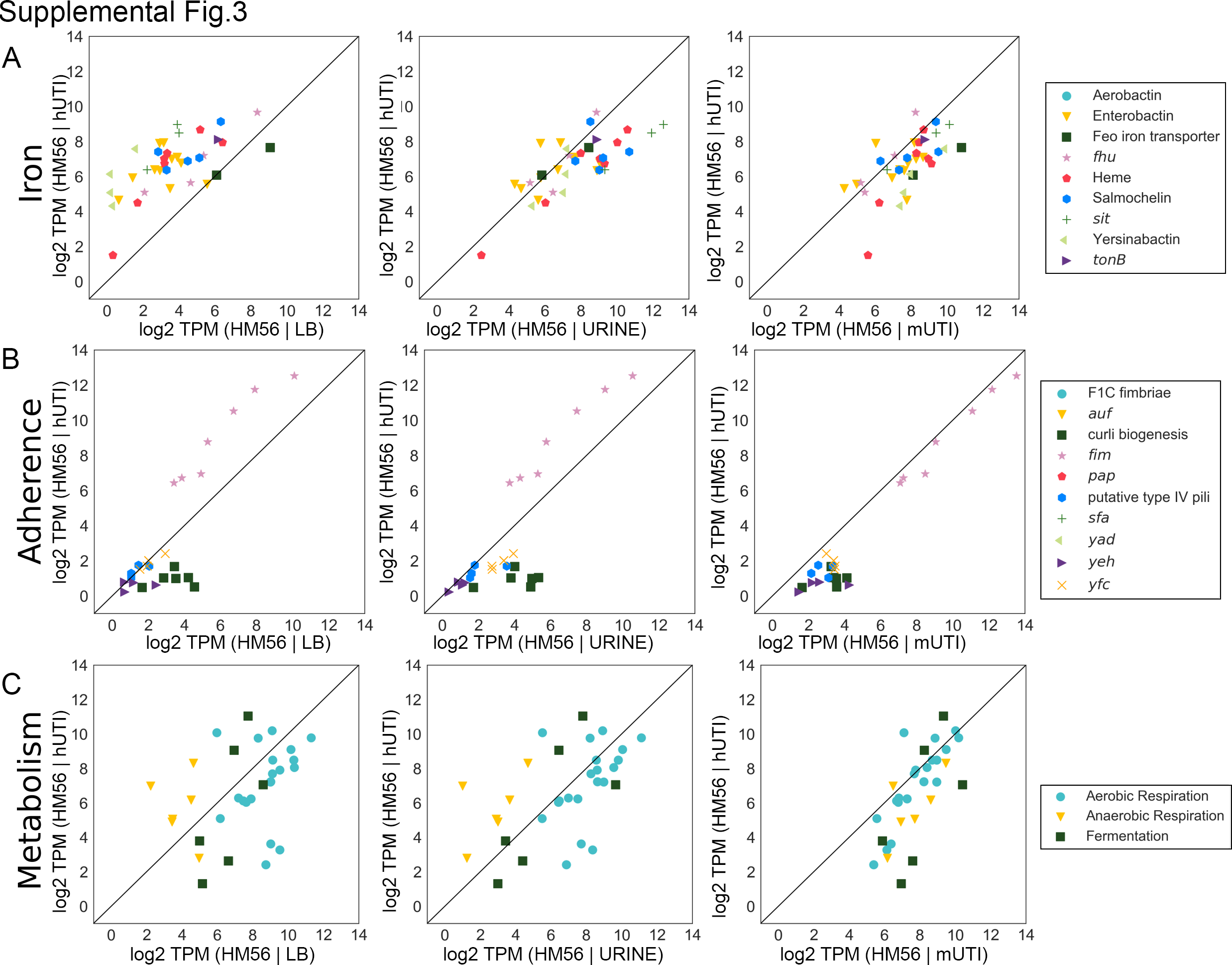

**Supplemental Figure 3. Gene expression of virulence factors as well as metabolic machinery is highly consistent between mouse model of UTI and human UTI in strain HM56.** Gene expression of iron acquisition operons (A), adherence genes (B), and metabolic pathways (C) for HM56 was compared between LB and human infection (LB vs hUTI), urine and human infection (urine vs hUTI), and mouse UTI and human UTI (mUTI vs hUTI).

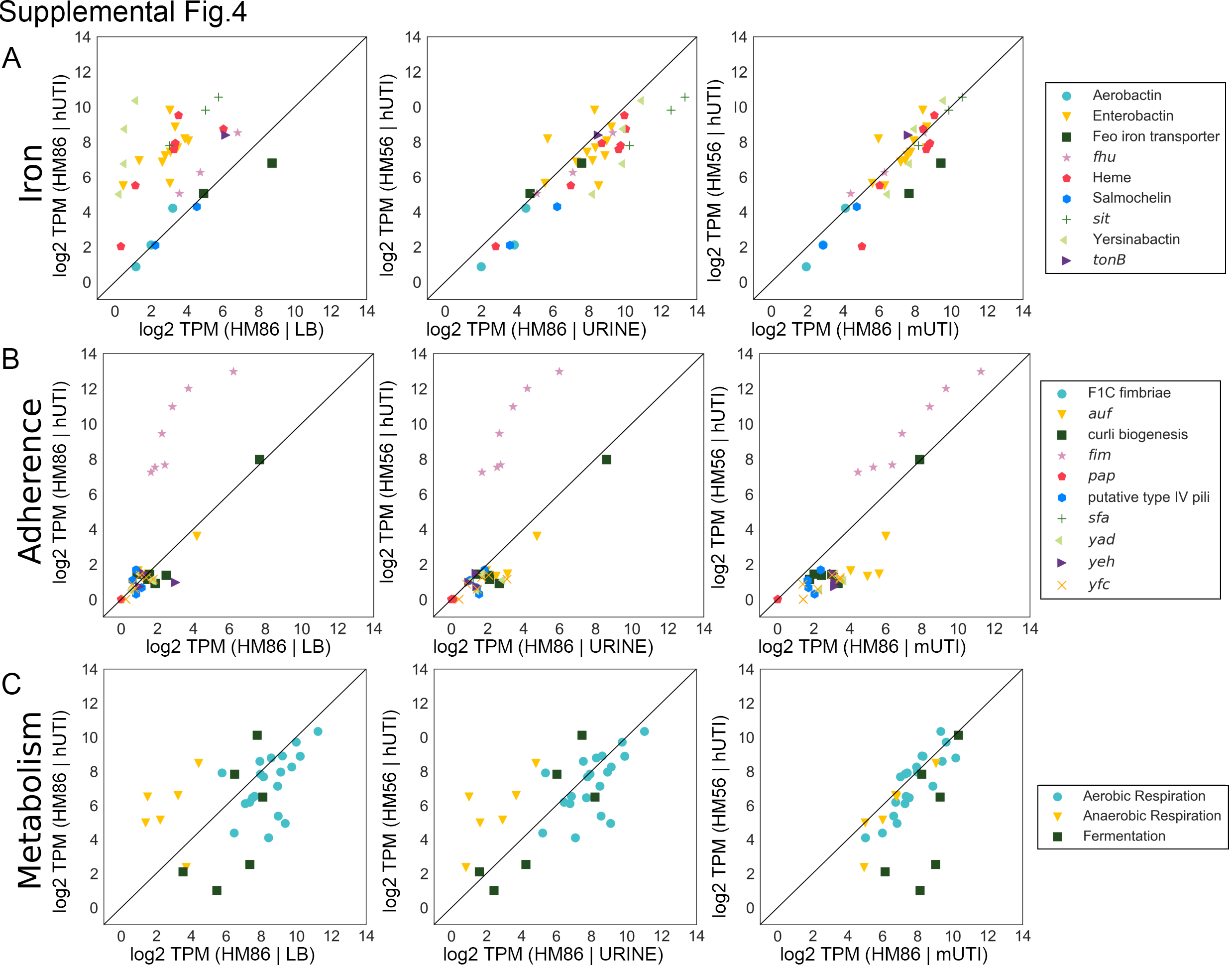

**Supplemental Figure 4. Gene expression of virulence factors as well as metabolic machinery is highly consistent between mouse model of UTI and human UTI strain HM86.** Gene expression of iron acquisition operons (A), adherence genes (B), and metabolic pathways (C) for HM86 was compared between LB and human infection (LB *vs* hUTI), urine and human infection (urine *vs* hUTI), and mouse UTI and human UTI (mUTI *vs* hUTI).
